# Engineering Stapled Peptide Inhibitors Reveals Design Principles for Targeting Talin-Induced Integrin Activation

**DOI:** 10.64898/2026.05.25.727761

**Authors:** Tong Gao, Elizaveta Belova, Yujiro Mori, Kathrine Wang, William H. Deni, Heinrich Roder, Bojana Gligorijevic, Jinhua Wu

**Affiliations:** Molecular Therapeutics Program, Fox Chase Cancer Center, Philadelphia, PA 19111, USA; Bioengineering Department, Temple University, Philadelphia, PA, USA; Archmere Academy, Claymont, DE, USA

## Abstract

Talin-induced integrin activation is a central regulator of cell adhesion and signaling. Intracellular targeting of this pathway remains challenging. Here, we report the structure-guided development of next-generation stapled peptidomimetics derived from the RIAM talin-binding site (TBS). Our structure-guided peptide optimization yielded S2-TBS and S3-TBS, resulting in talin binding and inhibitory potency, while improving thermal stability and cell uptake. NMR and crystallographic analyses confirmed conserved binding to talin head and talin rod domains. Functionally, optimized peptides potently inhibit talin-mediated integrin activation and suppress invadopodia-driven matrix degradation in cancer cells. To achieve more efficient and systematic peptide optimization beyond structure-guided design alone, we incorporated FoldX computational mutagenesis to quantitatively evaluate and prioritize favorable sequence substitutions across both talin-binding interfaces. Together, this work establishes an integrated structure- and computation-guided optimization strategy for the rational development of intracellular talin inhibitors.

**HIGHLIGHTS:**

1. S2-TBS increases binding affinity but introduces conformational heterogeneity due to a longer linker.
2. S3-TBS achieves improved stability, reduced size, and a well-defined stapled conformation.
3. Optimized peptides inhibit talin-mediated integrin function and suppress cancer cell matrix degradation.
4. FoldX-guided computational mutagenesis enables systematic optimization of talin-binding peptides.

## INTRODUCTION

Precise regulation of cell adhesion signaling is essential for hemostasis, immune responses, and cell-extracellular matrix communication.^1, 2^ This process depends largely on integrins, a family of cell receptors that transmit bidirectional signals across the plasma membrane (PM) to coordinate cellular functions including adhesion, migration, proliferation, and survival. ^3, 4^ Dysregulated integrin activation is implicated in thrombosis, immune dysfunction, inflammation, cardiovascular disease, and tumor metastasis. ^5, 6^ Integrin activation is largely governed by an inside-out signaling pathway in which adaptor proteins including talin and kindlin engage the β-integrin cytoplasmic tail, inducing conformational rearrangements of integrin ectodomains that promote ligand binding.^7^ Because talin serves as the key intracellular trigger of integrin activation, the talin:integrin interface represents an attractive target for selective therapeutic intervention.

Talin is a large cytoskeletal adaptor composed of an N-terminal FERM head domain containing four subdomains (F0, F1, F2, and F3) and an extended rod region containing multiple helical bundle domains (R1-R13) followed by a dimerization helix.^8–11^ Talin head domain (THD) interacts with the juxtamembrane (JM) helix and the NPxY motif of the integrin-β tail to induce integrin activation.^12–14^ In resting cells, talin adopts an autoinhibited conformation in which rod domains occlude the head domain.^15^ Talin activation is coordinated by RAP1 GTPase and its effector Rap1-interacting adaptor molecule (RIAM).^16, 17^ A short N-terminal α-helical segment of RIAM, termed the talin-binding site (TBS), binds both the F3 subdomain within THD and the talin rod region, thereby promoting talin activation and membrane translocation.^17–20^ Because the TBS-binding site on F3 lies adjacent to the integrin-β binding interface, a rationally engineered TBS-derived peptide could simultaneously interfere with talin:integrin-β association at the head domain and competitively block RIAM-mediated talin recruitment through the rod region.^21^ This dual targeting strategy, inspired by the intrinsic bispecificity of TBS toward talin head and rod domains, provides a “double-hit” mechanism for suppressing talin-mediated integrin activation.^21^

In our previous study, we developed a TBS-derived stapled peptidomimetic, S-TBS, that stabilizes the native α-helical conformation through a hydrocarbon linker.^21^ S-TBS exhibited enhanced helicity, improved cell permeability, and stronger affinity for talin, and effectively suppressed integrin activation in cells, establishing proof of concept for intracellular inhibition of integrin signaling by peptidomimetics. Structural analysis of the S-TBS:R7R8 complex further validated the design strategy by revealing that S-TBS engages the TBS-binding interface while positioning the hydrocarbon staple to sterically hinder THD:integrin-β interaction. Although S-TBS validated the feasibility of targeting talin using stapled peptidomimetics, further optimization was needed to improve binding potency, structural robustness, and overall molecular design for translational development. This unmet need motivated the development of next-generation analogs through structure-guided refinement of peptide sequence, length, and intrahelix staple.

Here, we report the rational optimization of next-generation talin-targeting stapled peptides through iterative structural, biochemical, and functional evaluation. Guided by the first-generation S-TBS scaffold, we systematically refined peptide sequence, length, and staple architecture to enhance talin engagement, as shown by increased affinity for both the talin head domain and rod domains, stronger inhibition of talin:integrin association, and more effective disruption of talin autoinhibition *in vitro*. NMR chemical shift perturbation analyses further confirmed that the optimized peptides retain binding to the canonical TBS-recognition site on the talin head while preserving the molecular basis for dual-site inhibition. Early optimization efforts improved biochemical potency but also exposed design tradeoffs, including conformational heterogeneity, limited crystal order, and reduced thermal stability associated with linker geometry and flexible peptide segments. These structural and biophysical insights guided subsequent refinement, yielding stapled peptides with improved helicity, enhanced thermal stability, well-defined molecular architecture at high resolution, and greater suppression of integrin-dependent cellular activities, including invadopodia-mediated matrix degradation. To further improve the efficiency of sequence optimization, systematic FoldX mutagenesis was used to evaluate substitutions across the TBS helix and identified several mutations predicted to enhance binding to both the talin head and rod domains. Together, these findings establish a structure- and computation-guided framework for iterative peptidomimetic optimization in which integrated structural, computational, and functional feedback can be leveraged to develop potent intracellular peptide inhibitors. More broadly, this work advances a promising therapeutic strategy for targeting integrin-driven pathologies and provides a generalizable blueprint for designing peptidomimetic modulators of complex intracellular signaling pathways.

## RESULTS

### Design of next-generation stapled peptides S2-TBS and S3-TBS

Guided by our previous characterization of the first-generation stapled peptide S-TBS, including structural insights from the crystal structure of the S-TBS:talin R7R8 complex (PDB: 7V1A), we sought to further optimize the TBS-derived stapled peptide scaffold through structure-guided refinement of sequence, length, and staple geometry.^21^ We have previously identified the T14E mutation within the TBS helix as an affinity-enhancing substitution for next-generation peptide designs (**Fig. 1A**).^21^ Based on the favorable helical stabilization and multi-site inhibitory mechanism established by S-TBS, we pursued two complementary optimization strategies: one aimed at extending the peptide scaffold and hydrocarbon linker to enhance talin engagement and steric interference with integrin-β binding, and the other focused on structural minimization by removing terminal residues that appeared disordered in the S-TBS:talin crystal structure while preserving the core talin-binding helix. One stapled peptide, termed S2-TBS, is derived from the TBS sequence of human RIAM residues 5-25 (Uniprot: Q7Z5R6). S2-TBS incorporates the T14E substitution with a C11 hydrocarbon staple linking residues 16 and 23 that are separated by two helical turns (residues *i, i+7*) in the TBS sequence (**Fig. 1B**).^21, 22^ This peptide was designed to test whether a more extended human sequence TBS would exhibit strong binding to talin, and whether a more extended staple would inhibit integrin-β binding to THD with higher potency. The other peptide, termed S3-TBS, contains human RIAM residues 7-25 with the T14E substitution. In contrast to the first-generation S-TBS peptide, which lacks the T14E mutation and contains additional N-terminal residues that are disordered in the S-TBS:R7R8 co-crystal structure, S3-TBS was rationally minimized to remove these structurally unresolved residues while preserving the core talin-binding helix (**Fig. 1B**).^21^ Together, these two design strategies enabled us to define how peptide architecture governs molecular recognition, structural stability, inhibitory activity, and molecular complexity, thereby guiding the development of more efficient and translationally favorable stapled inhibitors.

**Figure 1.**
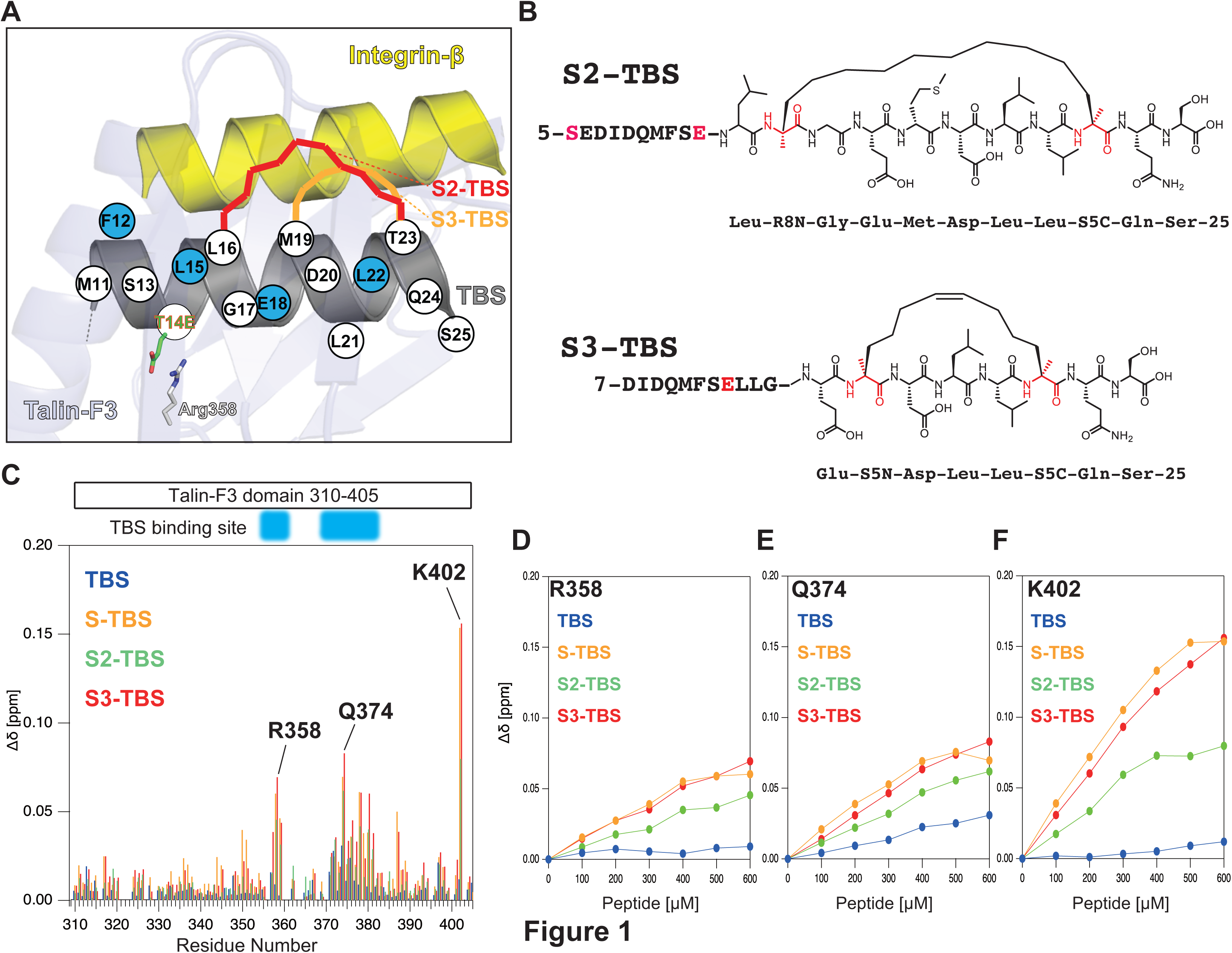
Structure-guided design and NMR characterization of stapled peptides. **(A)** Structure-guided design strategy for next-generation RIAM TBS-derived stapled peptides, highlighting optimization of sequence, length, and hydrocarbon staple geometry, including the T14E mutation. **(B)** Sequences and stapling configurations of WT-TBS, S-TBS, S2-TBS, and S3-TBS, illustrating differences in peptide length and staple type. **(C)** ^1^H - ^15^N HSQC spectra of ^15^N-labeled talin F2F3 in the presence of WT and stapled peptides. **(D-F)** Peptide concentration - dependent CSPs for representative residues within the talin F3 domain, including R358 **(D)**, Q374 **(E)**, and K402 **(F)**.

### NMR reveals conserved binding of next-generation stapled peptides to the talin head domain

Because crystallization of the talin head-stapled peptide complex was unsuccessful and direct visualization of the interaction could not be achieved, we performed 2D ^1^H-^15^N HSQC NMR spectroscopy using ^15^N-labeled talin F2-F3 tandem domains with wild-type TBS and stapled peptides (S-TBS^21^, S2-TBS, and S3-TBS). Consistent with the previously reported RIAM-TBS:Talin-F3 interface^17^, the wild-type TBS peptide induced distinct chemical shift perturbations (CSPs) at residues 357-360 and 370-380 (**Fig. 1C**). Importantly, all three stapled peptides produced highly similar CSP profiles, demonstrating that they retain engagement with the canonical TBS-binding surface on talin-F3. In particular, residues K357, G371, D372, Q374, D375, G376, and D397 exhibit comparable CSPs in WT-TBS and all stapled peptides, supporting a conserved binding interface on talin-F3 **(Supplemental Fig. 1)**. Notably, residue R358, which is expected to interact with S2- and S3-TBS via the engineered T14E mutation, exhibits strong CSPs during titration with S2- or S3-TBS peptides, consistent with the design rationale (**Fig. 1D**).^21^ Interestingly, S-TBS, which lacks T14E, also exhibits moderate perturbation at this position, suggesting that the hydrocarbon staple itself may alter local peptide positioning or broaden peptide engagement relative to WT-TBS.

Strong CSPs were also observed at residues Q374 and K402 in the presence of the stapled peptides. Q374, located within the β6-β7 loop of the F3 subdomain, also exhibits moderate but conserved CSPs with both WT and stapled peptides, suggesting a conserved interaction in the stapled peptides (**Fig. 1E**). In contrast, K402 exhibited minimal CSP with WT-TBS but strong, concentration-dependent shifts with stapled peptides (**Fig. 1F**). Because K402 contributes to stabilization of the active FERM configuration of talin, these enhanced perturbations suggest that stapled peptides engage an expanded surface extending toward the C-terminal region of F3.^23, 24^ Similarly, enhanced perturbation at W359, a residue involved in integrin- β recognition, further supports peptide interactions near the integrin-binding surface of THD, and is consistent with the inhibitory activity of the stapled peptides.^25, 26^

Additional minor CSPs were observed in distal regions of F2 and F3 (**Supplemental Fig. 1**). These effects likely reflect nonspecific interactions associated with the hydrophobic staple and the intrinsic flexibility of the isolated F2F3 construct that is structurally constrained in the context of the full-length talin head.^23^ Thus, these noncanonical perturbations are unlikely to represent physiologically relevant binding events. Together, these NMR analyses provide direct residue-level evidence that the next-generation stapled peptides engage the talin head domain through the canonical RIAM-interacting surface while exhibiting enhanced interactions with key THD residues involved in integrin activation.

### CD reveals improved conformational stability of next-generation stapled peptides

We evaluated the secondary structure and thermal stability of S-TBS, S2-TBS, and S3-TBS using circular dichroism (CD) spectroscopy. All three stapled peptides exhibited characteristic α-helical CD spectra with minima near 208 and 222 nm, consistent with successful stabilization of the helical TBS scaffold by hydrocarbon stapling (**Fig. 2A**).^27^ Among the peptides, S3-TBS displayed the strongest ellipticity at 222 nm, suggesting enhanced helical stabilization and reduced conformational flexibility relative to S2-TBS. This observation is consistent with the more compact peptide architecture and the shorter (*i, i+4*) hydrocarbon staple incorporated in S3-TBS.

**Figure 2.**
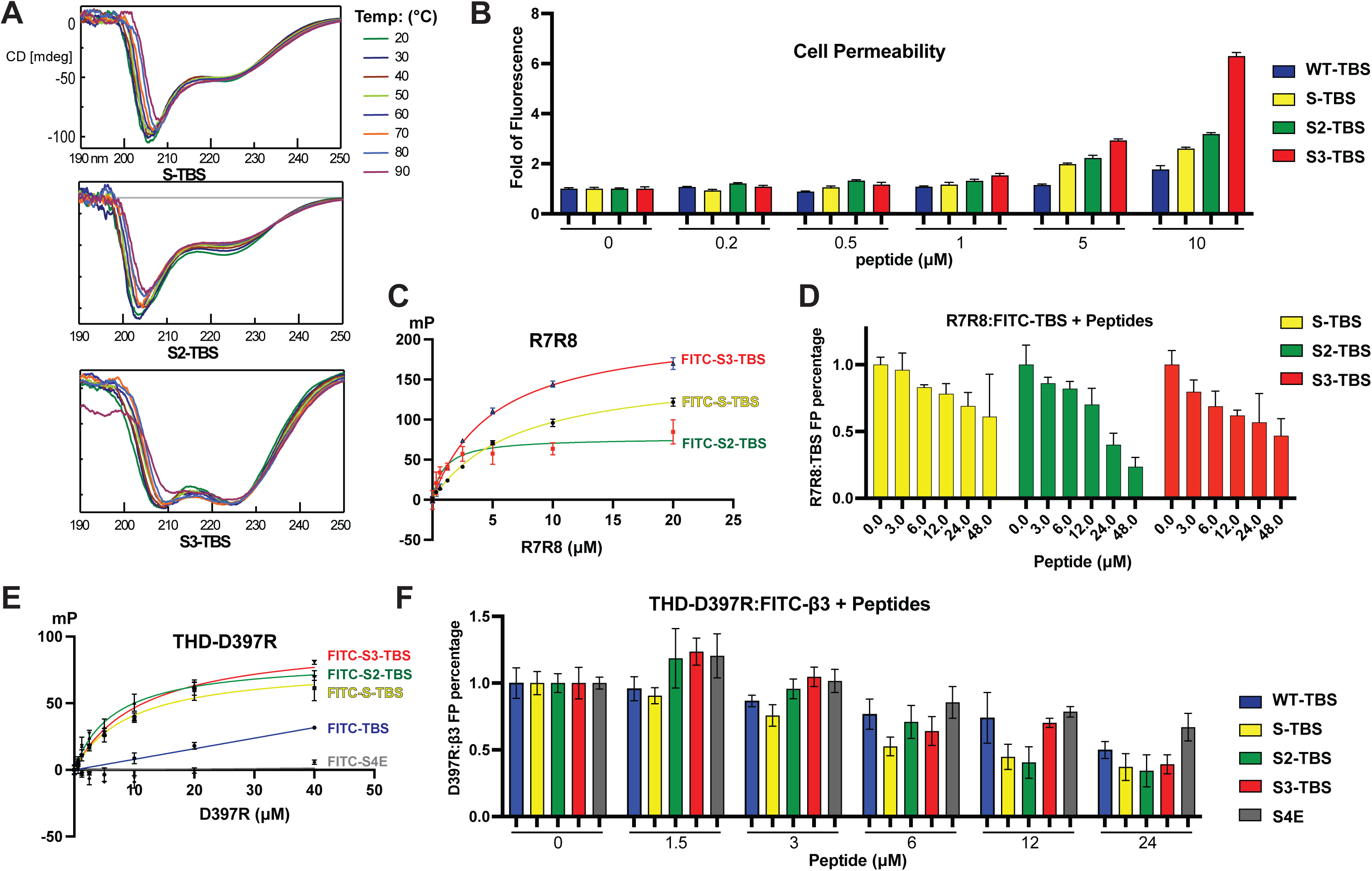
Biophysical and functional characterization of stapled peptides. **(A)** CD spectra at 20°C demonstrating α-helical structure. All peptides exhibited α-helical characteristics, with S3-TBS showing enhanced ellipticity and improved thermal stability. **(B)** Cellular permeability of WT-TBS and stapled peptides in CHO cells, showing enhanced uptake for all stapled peptides, with S3-TBS exhibiting the highest accumulation. **(C)** Temperature-dependent CD showing retention of helicity, with improved stability for S3-TBS. **(D)** FP binding to talin R7R8, showing enhanced binding of stapled peptides. **(E)** FP binding to THD-D397R, indicating increased interaction with the talin head. **(F)** Competitive FP assay showing dose-dependent inhibition of THD:β3 interaction, with S2-TBS exhibiting the strongest inhibition at intermediate concentrations. Data are mean ± SD. THD-D397R enhances integrin-binding sensitivity.

To further assess thermal stability, CD spectra were collected from 20°C to 90°C at 10°C intervals. All stapled peptides retained their α-helical spectral features across the examined temperature, although gradual reductions in ellipticity were observed at elevated temperatures (**Fig. 2A**). Notably, S3-TBS maintained comparatively stronger ellipticity at 222 nm throughout the temperature scan, indicating improved helical stability relative to S-TBS and S2-TBS.^28^ Together, these data support that structural minimization and optimization of staple geometry in S3-TBS enhance conformational stability of the stapled peptide scaffold.

### S3-TBS exhibits enhanced cell permeability

To further evaluate the biochemical and functional properties of the next-generation stapled peptides, we assessed their cellular permeability. Compared with the WT-TBS peptide, all stapled peptides exhibited substantially enhanced cellular uptake in CHO cells, consistent with stabilization of the helical peptide scaffold by hydrocarbon stapling. Among the tested peptides, S3-TBS displayed the highest permeability, exhibiting markedly increased intracellular accumulation at 5-10 μM relative to S-TBS and S2-TBS. In contrast, WT-TBS exhibited minimal uptake except at the highest concentrations tested (**Fig. 2B**). Interestingly, although S-TBS and S2-TBS share similar peptide lengths but differ in hydrocarbon staple, the longer and more flexible C11 staple in S2-TBS did not substantially impair the cell permeability relative to the C8-stapled S-TBS. In contrast, S3-TBS combines a shorter C8 staple geometry with a minimized peptide sequence, resulting in the most compact overall molecular architecture among the stapled peptides tested. S3-TBS exhibits significantly higher cellular uptake than S-TBS and S2-TBS. These findings suggest that overall peptide size and sequence length contribute more significantly to cellular permeability than moderate changes in staple length, and the more flexible C11 staple remains generally well tolerated in terms of membrane permeability. The enhanced permeability of S3-TBS supports that structural minimization together with a compact hydrocarbon staple can improve intracellular delivery of stapled peptide inhibitors.

### Next-generation stapled peptides exhibit enhanced affinities with both talin head and talin rod domains

We next characterized the interactions of the stapled peptides with talin rod R7R8 domains and the THD using fluorescence polarization (FP) assays. All stapled peptides bound talin R7R8 with substantially greater affinity than the WT-TBS peptide (**Fig. 2C**). While S3-TBS (*K_d_* = 5.1 μM) exhibits enhanced binding to R7R8 than S-TBS (*K_d_* = 7.0 μM), S2-TBS exhibits the strongest binding affinity toward R7R8 with a *K_d_* of 0.98 μM (**Table 1**). To determine whether the next-generation peptides inhibit talin-mediated interactions, we performed competitive FP assays using the R7R8:FITC-TBS complex. All stapled peptides inhibited the R7R8:TBS interaction in a dose-dependent manner (**Fig. 2D**). Consistent with the binding data, while both S2-TBS and S3-TBS exhibit stronger inhibition of the TBS:R7R8 interaction than S-TBS, S2-TBS exhibits the strongest inhibition (**Supplemental table 1**). Notably, S2-TBS reduced R7R8:TBS association to ∼40% and ∼25% at 24 and 48 μM, respectively, whereas S3-TBS exhibited intermediate inhibition and S-TBS exhibited more modest inhibition. These findings suggest that the extended peptide sequence and C11 staple of S2-TBS enhance talin rod engagement and disrupt RIAM-mediated talin recruitment more effectively.

**Table 1.**
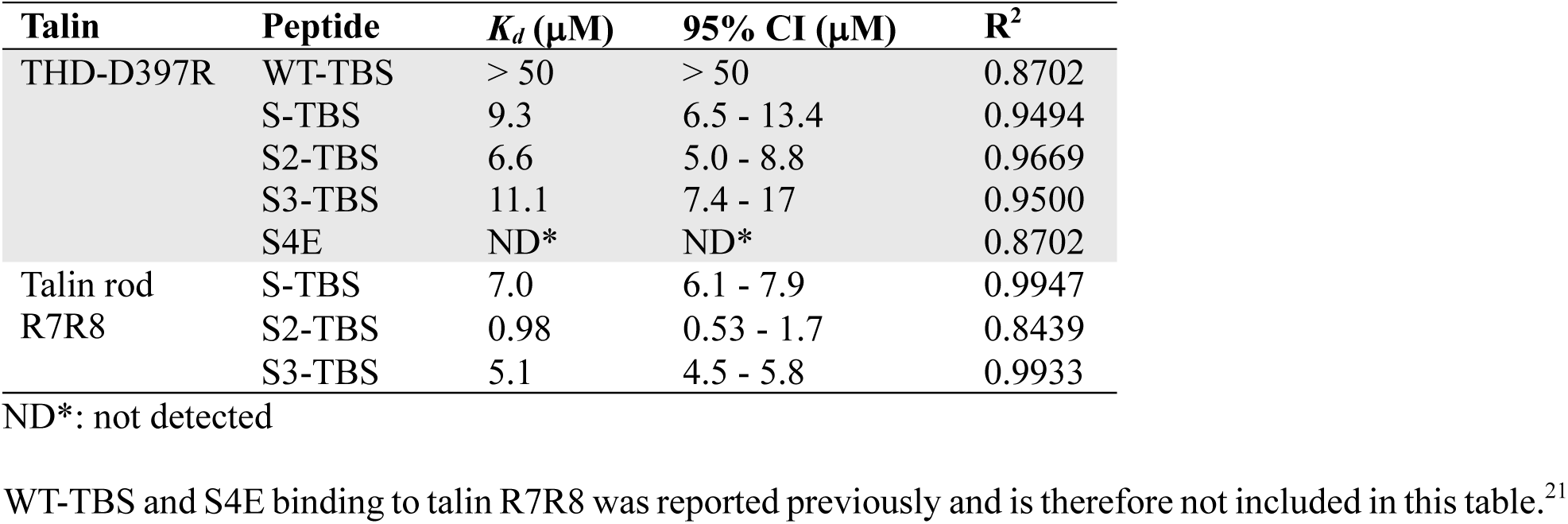
Binding affinities measured by fluorescence polarization.

To better evaluate peptide interactions with THD, we used a THD-D397R mutation, which exhibits enhanced integrin binding by stabilizing the C-α helix within the F3 domain. WT-TBS and a non-interacting S4E (S-TBS with M11E/F12E/L15E/L16E substitutions) control peptide were also included to validate the binding specificity of the peptides with THD-D397R.^23^ All stapled peptides demonstrated enhanced interactions with THD-D397R compared with WT-TBS (**Fig. 2E**), consistent with the stronger CSPs observed in the NMR analyses. No interaction was detected for the S4E peptide. Notably, THD-D397R exhibits stronger interaction with S-TBS (*K_d_* = 9.3 μM) than wild-type THD (*K_d_* = 38 μM). We next evaluated inhibition of the talin:integrin interaction using a THD-D397R:FITC-β3 competitive FP assay (**Fig. 2F, Supplemental Table. 1**). Consistent with the original design rationale of sterically interfering with the integrin-binding surface of talin-F3, all stapled peptides inhibited talin:β3 association in a concentration-dependent manner. S-TBS, S2-TBS, and S3-TBS all exhibited substantially stronger inhibition than WT-TBS at intermediate peptide concentrations. S2-TBS exhibits the strongest inhibition of the THD:β3 interaction at 12 μM and 24 μM, suggesting that the larger C11 staple in S2-TBS enhances steric interference with the integrin-binding interface on THD, resulting in more potent inhibition than the C8-stapled S-TBS and S3-TBS peptides. Together, these results demonstrate that the optimized stapled peptides preserve strong talin binding while exhibiting enhanced inhibitory effects on both talin rod and talin:integrin interactions.

### Crystal Structures of S2-TBS and S3-TBS in complex with talin R7R8

To directly examine how the next-generation stapled peptides interact with talin and how modifications in staple geometry and peptide architecture influence structural organization, we determined crystal structures of S2-TBS and S3-TBS in complex with talin R7R8 domains. Crystals of the S2-TBS:R7R8 complex diffracted to 3.0-Å resolution **(Table 2)**, revealing a binding configuration similar to that of the S-TBS:R7R8 complex (**Fig. 3A**). However, despite preserving the overall binding orientation, the S2-TBS complex exhibited weaker electron density surrounding the hydrocarbon linker and elevated overall disorder. In particular, the extended C11 staple was only partially resolved, suggesting considerable conformational flexibility within the staple region (**Fig. 3B**). This structural heterogeneity is consistent with the poorer diffraction quality and elevated B-factors observed for the S2-TBS complex. Nonetheless, surface analysis of the S2-TBS:R7R8 interface demonstrated that the peptide maintains extensive hydrophobic interactions with talin-R8 despite the partially disordered linker (**Fig. 3C**). The extended C11 staple occupies the same surface region as the C8 staple in S-TBS but appears more dynamic, suggesting that the longer linker preserves talin engagement while increasing both binding surface coverage and conformational flexibility. These observations thus support the strong inhibitory activity of S2-TBS toward THD:β3 interaction despite its reduced structural rigidity.

**Figure 3.**
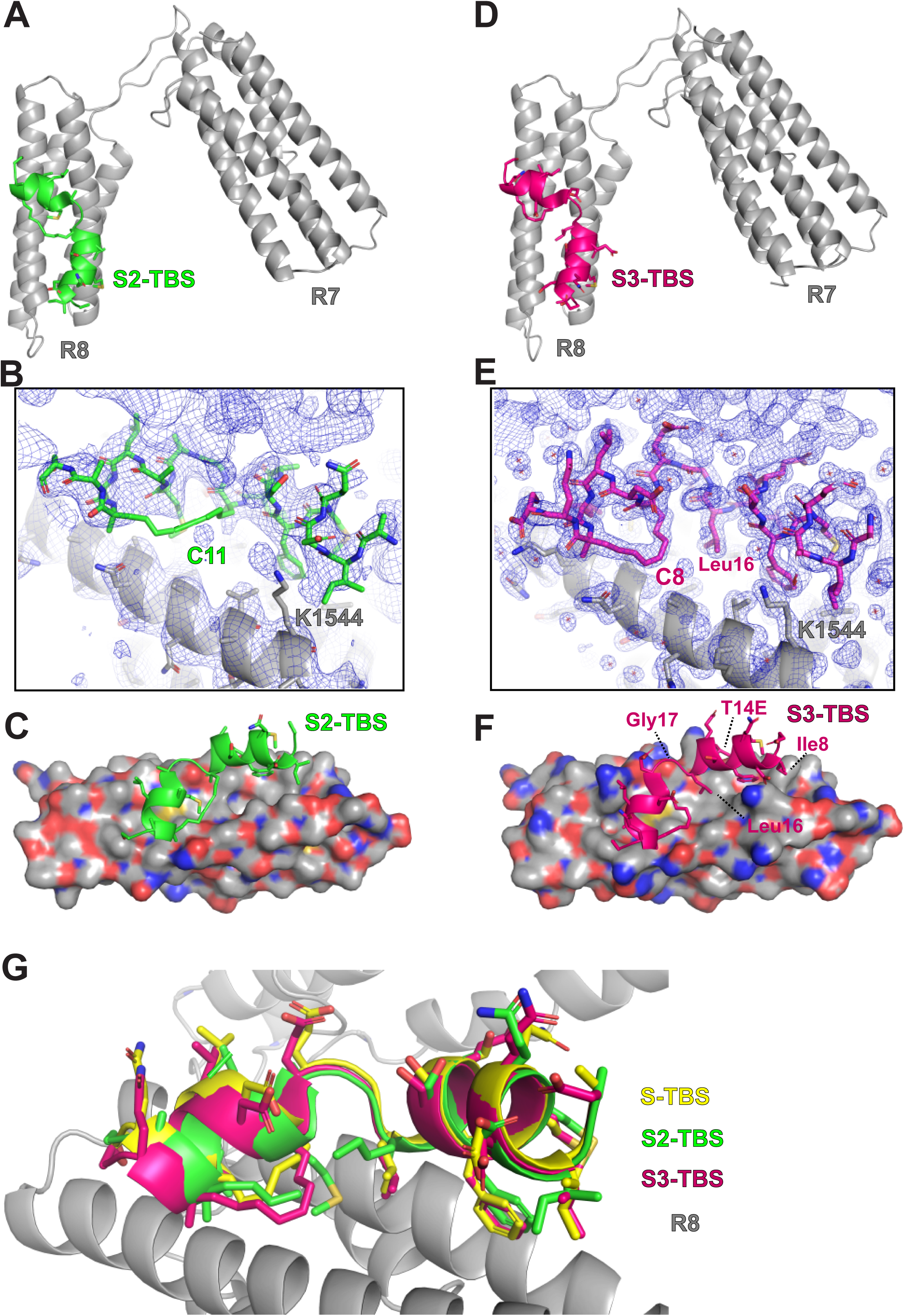
Structural basis of stapled peptide interaction with talin R7R8. **(A)** Overall structure of S2-TBS bound to R7R8. **(B)** 2Fo-Fc electron density at the S2-TBS:R7R8 interface. (**C)** Surface interactions with R7R8 in surface representation and S2-TBS in cartoon representation. **(D)** Overall structure of S3-TBS bound to R7R8. **(E)** 2Fo-Fc electron density at the S3-TBS:R7R8 interface. (**F)** Surface interactions with R7R8 in surface representation and S3-TBS in cartoon representation. The well-ordered Leu16 is shown in **(E)** and **(F)** to highlight the partially occupied pocket on R7R8. **(G)** Superposition of S-TBS, S2-TBS, and S3-TBS demonstrating conserved binding modes.

**Table 2.** Data collection and refinement statistics.

| **Crystals** | **S2-TBS:R7R8** | **S3-TBS:R7R8** |
| --- | --- | --- |
| **PDB Entry** | **37TO** | **37TQ** |
| **Data Collection** |  |  |
| **Light Source** | NSLS-II AMX | NSLS-II AMX |
| **Wavelength (Å)** | 0.92010 | 0.92010 |
| **Resolution range (Å)** | 40.0 - 3.0 (3.44 - 3.0) | 29 - 1.68 (1.72 - 1.68) |
| **Space group** | P2<sub>1</sub>2<sub>1</sub>2 | P2<sub>1</sub>2<sub>1</sub>2 |
| **Unit cell** |  |  |
| **a, b, c (Å)** | 60.3, 105.6, 49.2 | 59.7, 107.0, 48.8 |
| **α, β, γ (°)** | 90, 90, 90 | 90, 90, 90 |
| **Multiplicity** | 12.5 (13.4) | 9.3 (7.5) |
| **Completeness (%)** | 100 (100) | 100 (100) |
| **Mean I/sigma(I)** | 11.6 (0.9) | 9.2 (0.9) |
| **R-merge** | 0.137 (3.3) | 0.128 (2.6) |
| **CC(1/2)** | 0.998 (0.44) | 0.998 (0.393) |
| **No. Reflections** | 6668 (2162) | 36588 (2718) |
| **Refinement** |  |  |
| **R-work** | 0.2519 (0.3545) | 0.2073 (0.3275) |
| **R-free** | 0.3174 (0.4311) | 0.2375 (0.3601) |
| **Number of atoms** |  |  |
| **macromolecules** | 2275 | 2387 |
| **ligands** | 0 | 0 |
| **solvent** | 3 | 206 |
| **RMSD bonds(Å)** | 0.003 | 0.008 |
| **RMSD angles (°)** | 0.63 | 0.91 |
| **Average B-factor (Å<sup>2</sup>)** | 112.90 | 35.67 |
| **Macromolecules (Å<sup>2</sup>)** | 112.93 | 34.99 |

Statistics for the highest-resolution shell are shown in parentheses.

We next determined the structure of S3-TBS bound to talin R7R8. Similar to S2-TBS, S3-TBS occupies the canonical TBS-binding groove on R8 and adopts a conserved α-helical configuration (**Fig. 3D**). In contrast to S2-TBS, the S3-TBS complex exhibited highly ordered electron density throughout the peptide and hydrocarbon staple at 1.68-Å resolution **(Table 2)**. The C8 linker was fully resolved with well-defined electron density (**Fig. 3E**), supporting a substantially more rigid and conformationally restrained architecture. Correspondingly, the S3-TBS:R7R8 structure displayed markedly improved crystallographic quality and significantly lower B-factors relative to the S2-TBS complex. Surface representation of the S3-TBS:R7R8 interface further demonstrated that sequence minimization does not compromise talin recognition (**Fig. 3F**). Despite removal of terminal disordered residues in S-TBS:R7R8 complex, S3-TBS preserves the key hydrophobic interactions essential for talin binding with a more compact and ordered molecular surface. These findings are consistent with the enhanced helicity, thermal stability, and cellular permeability observed for S3-TBS.

Superposition of S-TBS, S2-TBS, and S3-TBS bound to R7R8 revealed highly conserved peptide configurations and overall binding modes, indicating that both peptide truncation and staple modification preserve recognition of the talin-R8 interface (**Fig. 3G**). Nevertheless, local structural differences were observed around the staple region. While the C8 staple of S-TBS and S3-TBS adopts a relatively constrained orientation against the talin surface, the C11 linker in S2-TBS exhibits substantially greater conformational flexibility, suggesting that the staple length strongly affects molecular order without abolishing target binding.

### S2-TBS and S3-TBS inhibits talin-induced integrin activation

We next examined whether the optimized stapled peptides suppress talin-induced integrin activation in cells using a fluorescence-activated cell sorting assay.^29, 30^ CHO-A5 cells, which stably express integrin αIIbβ3, were transfected with GFP-tagged talin head domain (GFP-THD) and Kindlin-2, and integrin activation was monitored by an activation-specific antibody PAC-1. As observed previously, S-TBS inhibited talin-induced integrin activation at both 5 and 25 μM (**Fig. 4A**).^21^ Both S2-TBS and S3-TBS also inhibited talin-induced integrin activation to levels near that ovserved with the inactive talin W359A mutant at both concentrations (**Fig. 4A**), although their inhibitory effects were not significantly different at two tested concentrations. Nevertheless, these results further demonstrate that new generation stapled peptides can suppress talin-dependent integrin activation in cells, while suggesting that this assay may be less sensitive for distinguishing differences in activity among the three peptides. We therefore further compared their cellular activities using an invadopodia-mediated matrix degradation assay in BT-549 cells.

**Figure 4.**
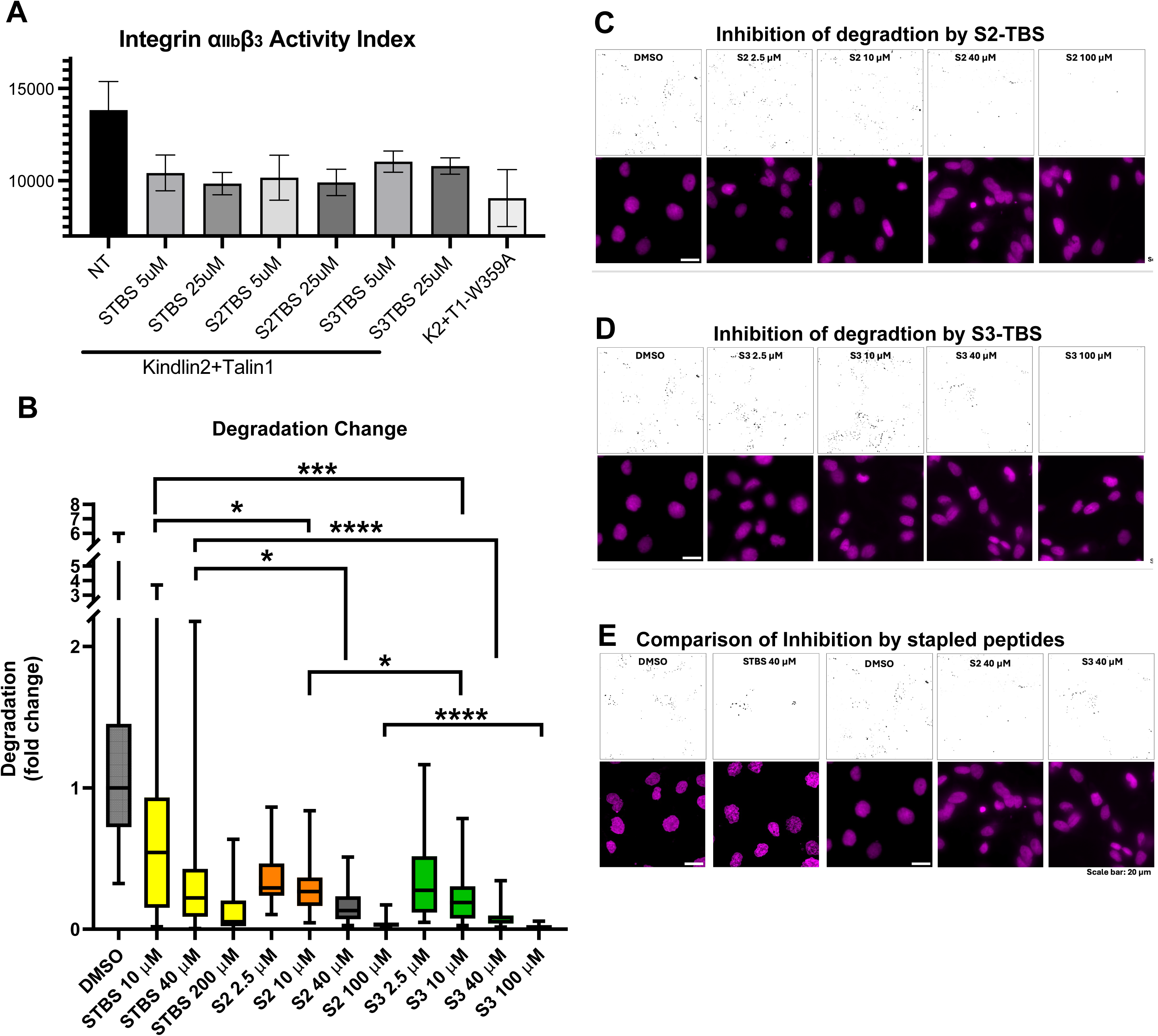
Inhibition of invadopodia activity in BT-549 cells by stapled peptides. **(A)** S-TBS, S2-TBS, and S3-TBS inhibit integrin αIIbβ3 activation. CHO-A5 cells were co-transfected with GFP-talin1 and kindlin2 and treated with the indicated peptides. Integrin activation was measured by FACS using PAC-1 and expressed as the αIIbβ3 activity index. **(B)** Quantification of invadopodia-mediated gelatin degradation, expressed as fold change relative to the DMSO-treated condition. **(C)** Representative images of gelatin degradation by BT-549 cells treated with increasing concentrations of the S2 peptidomimetic. Nuclei are shown in magenta (DRAQ5). **(D)** Representative images of gelatin degradation by BT-549 cells treated with increasing concentrations of the S3 peptidomimetic. **(E)** Representative images showing gelatin degradation (black dots) by BT-549 cells treated with different peptidomimetics at a common concentration of 40 μM. Data represent two biological replicates with two technical replicates each. Statistical significance was determined using a two-tailed Mann-Whitney test (*P* < 0.05, *; *P* < 0.01, **; *P* < 0.001, ***; *P* < 0.0001, ****; ns, not significant). Scale bar = 20 μm.

### S2-TBS and S3-TBS inhibit invadopodia-mediated matrix degradation more strongly in BT-549 cells

Because talin-mediated integrin signaling is required for invadopodia formation and extracellular matrix remodeling during tumor invasion, we next evaluated whether the optimized stapled peptides suppress invadopodia-mediated matrix degradation in BT-549 breast cancer cells.^31–33^ BT-549 cells were plated on fluorescently labeled gelatin and treated with different concentrations of peptidomimetics (**Fig. 4B**).. Representative images show areas of fluorescently labeled gelatin degradation (black dots) and cell nuclei (magenta, DRAQ5) under various treatment conditionsh.

Representative images revealed dose-dependent inhibition of matrix degradation by both S2-TBS and S3-TBS. Increasing concentrations of S2-TBS (2.5 - 100 μM) progressively reduced gelatin degradation, with substantial suppression observed at 40 and 100 μM (**Fig. 4C**). A similar concentration-dependent effect was observed for S3-TBS, which also markedly inhibited matrix degradation at higher concentrations (**Fig. 4D**). To directly compare inhibitory potency among stapled peptides, BT-549 cells were treated with S-TBS, S2-TBS, or S3-TBS at a common concentration of 40 μM (**Fig. 4E**). All three peptides substantially reduced matrix degradation relative to the DMSO control. Consistent with the enhanced biochemical and structural properties observed in earlier analyses, S2-TBS and S3-TBS produced stronger suppression of matrix degradation than the first-generation S-TBS peptide.

Quantification of degradation area confirmed these observations (**Fig. 4B**). All stapled peptides significantly reduced matrix degradation compared with the DMSO-treated cells, demonstrating effective inhibition of invadopodia activity.^33^ Consistent with their enhanced talin-binding and inhibitory properties, both S2-TBS and S3-TBS exhibited greater potency than the first-generation S-TBS peptide. Interestingly, although S2-TBS displayed the strongest talin binding and inhibition in biochemical assays, S3-TBS exhibited slightly stronger suppression of matrix degradation at higher concentrations (40-100 μM). This enhanced cellular activity is consistent with earlier findings demonstrating improved conformational stability and superior cellular permeability of S3-TBS. Together, these results suggest that although strong talin engagement contributes to inhibitory potency, optimized molecular compactness and intracellular delivery can further enhance functional efficacy in cellular settings.

### Systematic computational mutagenesis identifies experimentally testable mutations for improving talin engagement

The structures of R7R8 in complex with the stapled peptides revealed an extended TBS-binding surface, suggesting that several peptide residues may provide opportunities for further optimization. To systematically evaluate the contribution of each residue within the TBS helix, we performed computational mutagenesis using FoldX across the core TBS sequence from Asp7 to Ser25.^34^ Each residue was substituted with the other 19 amino acids, and the resulting 361 substitutions were analyzed for binding free energy (ΔΔG) using all 20 NMR conformers for THD:TBS (PDB: 2MWN) and one crystallographic R7R8:TBS (PDB: 4W8P) structural models (**Supplemental Table 1**). For each substitution, the lowest ΔΔG value among the 20 NMR conformers of THD:TBS was used for assessment. Among all substitutions, 75 were predicted to stabilize both complexes, 85 stabilized only THD, 52 stabilized only R7R8, and 149 destabilized both interfaces.

FoldX predictions were highly consistent with our previous experimental data for mutations that impair talin binding. Most substitutions at residues directly contacting R7R8, including Met11, Phe12, Leu15, Glu18, Met19, Leu21, and Leu22, were predicted to be unfavorable (**Fig. 5A**), underscoring the critical roles of these residues in maintaining the R7R8-binding interface. Consistent with these predictions, we have previously showed that the L15Y, L15W, and E18A mutations all markedly reduced R7R8 binding, providing experimental validation of the FoldX analyses (**Fig. 5B**).^18^ Similarly, S13G, which disrupts the kink in the TBS helix and abolished R7R8 binding in the pulldown assay, was also unfavored in the FoldX analysis.^18^ Thus, the computational analysis showed particularly strong predictive power for identifying substitutions that disrupt the talin interaction.

**Figure 5.**
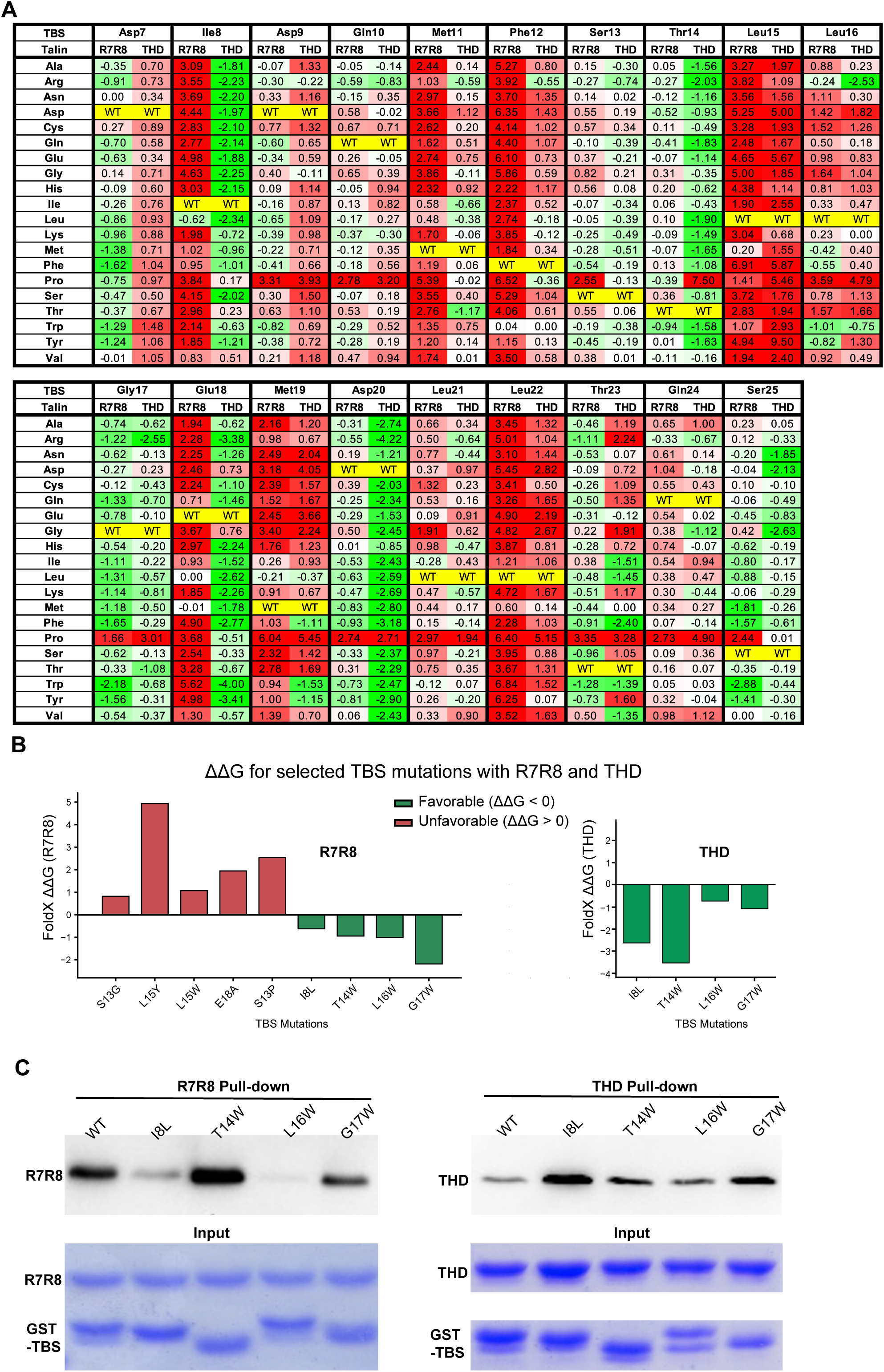
Computational mutagenesis and experimental validation of TBS mutations for talin binding. (A) Systematic FoldX mutagenesis of RIAM TBS residues for binding to talin R7R8 and THD. Each residue was substituted with the other 19 amino acids, and the predicted changes in binding free energy (ΔΔG) were calculated using 20 NMR conformers for each structural model. The lowest ΔΔG value among the 20 conformers was used for each substitution. Green indicates favorable substitutions (ΔΔG < 0), whereas red indicates unfavorable substitutions (ΔΔG > 0). WT residues are indicated in yellow. **(B)** FoldX-predicted ΔΔG values for selected TBS mutations. Previously characterized mutations with unfavorable effects on R7R8 binding (S13G, L15Y, L15W, E18A, and S13P) are shown together with four substitutions (I8L, T14W, L16W, and G17W) predicted to enhance binding. Predicted ΔΔG values for the four favorable substitutions with THD are shown separately. **(C)** GST-TBS pull-down of talin R7R8 or THD. GST-tagged WT or mutant TBS peptides, I8L, T14W, L16W, and G17W, were incubated with purified talin R7R8 (left) or THD (right) with a His6 tag, and bound proteins were analyzed by Western blot.

We then examined whether the same analysis could be used to identify substitutions that enhance talin binding. Among the favorable substitutions, I8L, T14W, L16W, and G17W were selected because they were predicted to improve binding to both R7R8 and THD (**Fig. 5B**). To test these four substitutions for binding to talin, we performed GST-TBS pull-down assays with purified R7R8 and THD (**Fig. 5C**). Several of the selected substitutions showed enhanced binding to either R7R8 or THD in the pull-down assays. These results support the utility of the FoldX analysis for prioritizing substitutions with the potential to improve talin engagement. Nevertheless, several of the predicted substitutions produced enhanced binding in a domain-dependent manner. T14W showed a clear increase in R7R8 binding compared with WT TBS, whereas I8L and G17W showed improved binding to THD. These results show that FoldX can identify candidate substitutions with gain-of-binding potential and help prioritize mutations for experimental testing. This systematic computational screening approach significantly narrows the candidate range by filtering out unfavorable substitutions and focusing experimental efforts on a small set of favorable candidates.

Together, these results define both the strengths and limitations of systematic computational mutagenesis for peptide optimization. FoldX showed excellent agreement with experimentally characterized loss-of-function mutations and therefore provides a reliable and efficient means of eliminating unfavorable substitutions. Although prediction of gain-of-function mutations was less direct, computational screening narrowed the search to a small number of candidates for experimentally testing and successfully identified substitutions that enhanced binding to either R7R8 or THD. Integration of this computational screening with biochemical validation therefore provides a practical strategy for identifying and refining sequence modifications for the next generation of TBS-based stapled peptides.

## DISCUSSION

In this study, we employed two complementary design strategies to optimize RIAM TBS-derived stapled peptides targeting talin. S2-TBS was designed by extending both the peptide sequence and hydrocarbon staple, with the goal of increasing talin engagement and enhancing steric interference with talin-mediated integrin binding. In contrast, S3-TBS was designed through structural minimization, removing disordered terminal residues while preserving the core talin-binding helix and maintaining the shorter C8 staple for conformational control. The results reveal an important tradeoff in stapled peptide optimization. S2-TBS exhibited strong biochemical activity, particularly in binding to talin R7R8 and THD, as well as disrupting the talin R7R8:TBS interaction, supporting the idea that an extended peptide and longer staple can enhance target engagement and inhibitory potency. However, the longer C11 staple in S2-TBS also appeared to introduce conformational flexibility, as indicated by the poor structural ordering of the staple, and suboptimal cell-based performance compared with S3-TBS. This suggests that increasing molecular size or linker length may improve *in vitro* binding properties but can compromise conformational stability, permeability, or intracellular efficacy that are essential for effective intracellular inhibition. ^35^ ^36, 37^

By contrast, S3-TBS demonstrates the advantages of a more compact and structurally constrained scaffold. S3-TBS retained the canonical talin-binding mode, showed improved helicity and thermal stability, exhibited stronger cellular permeability and potent inhibition in cell-based matrix degradation assays, likely benefiting from its shorter sequence and more closely positioned staple.^38–40^ The high-resolution S3-TBS:R7R8 structure further supported that the shorter C8 staple is well ordered and compatible with the talin-binding surface. These findings suggest that structure-guided peptide minimization can improve molecular stability and functional efficacy without compromising target recognition. Thus, an optimal intracellular stapled peptide inhibitor require a balance between strong target engagement and compact molecular architecture.

Systematic computational mutagenesis provides a practical and reliable optimization strategy for the development of peptidomimetic inhibitors. Our FoldX results were in excellent agreement with the previous experimental mutagenesis data and supported the T14E substitution incorporated into S2-TBS and S3-TBS. More importantly, the analysis identified several previously unrecognized residues that may provide additional improvements in talin binding. Several classes of favorable substitutions emerged from the analysis. In particular, Gly17 displayed an unexpectedly high degree of mutational tolerance, with 17 of 19 substitutions predicted to stabilize both THD and R7R8 interactions. Because Glycine has a relatively low helical propensity, these findings suggest that enhancing the helical stability of TBS may improve talin engagement. In contrast, L16W, despite favorable FoldX predictions for both complexes, did not improve binding in the pull-down assays and substantially reduced R7R8 binding. This discrepancy emphasizes that calculated binding energies capture only part of the energetic and conformational requirements governing peptide-protein interactions.These findings demonstrate that systematic computational mutagenesis can effectively prioritize sequence positions for experimental validation and facilitate the rational optimization of peptidomimetic inhibitors.

A future direction will be to integrate the experimental and computational approaches as we show here. Sequence features that enhance binding, such as those seen in S2-TBS or predicted by computational mutagenesis, can be incorporated into a more compact S3-TBS-like scaffold, while maintaining the shorter and well-ordered staple geometry. This approach may allow further improvement of potency without compromising permeability or structural stability. With recent advances in AI-driven molecular modeling, this integrated experimental-computational approach will undoubtedly accelerate the rational development of stapled peptides and other peptidomimetic inhibitors targeting challenging intracellular protein-protein interactions.

## AUTHOR CONTRIBUTIONS

T.G. purified proteins, performed crystallization experiments, and collected the crystallographic data. T.G. and J.W. determined the structures. Y.M. performed NMR analyses. W.H.D. performed CD analyses. T.G. performed cell permeability, FP binding, FACS, and competition assays. E.B. performed the invadopodia degradation assays. K.W. and J.W. performed the FoldX analysis. H.R. and B.G. contributed to the data analyses. J.W. supervised the project and served as the principal manuscript author.

## ACKNOWLEDGMENTS

We thank the beamline staff at AMX and FMX of the National Synchrotron Light Source-II, Brookhaven National Laboratory, and beamline 7B2 at MacCHESS, Cornell University, for technical support. This work was supported by an NIH Grant GM119560 (to J.W.), an ASH bridge grant (to J.W.), and Pennsylvania Department of Health Grant 4100085739 (to J.W.). T.G. was also supported by the Elizabeth Knight Patterson Postdoctoral Fellowship.

## EXPERIMENTAL PROCEDURES

### Plasmid construction and protein purification

Talin R7R8 and talin head were subcloned into a modified pET28a vector with a His6-tag in previous studies. ^21, 41^ They were expressed and purified as described previously. ^21, 24, 42^ Briefly, *E. coli* BL21(DE3) transformed with modified pET28a-talin constructions were cultured in fresh LB medium containing 50 μg/ml kanamycin with shaking flasks at 200rpm, 37℃. When the OD at 600nm reached 0.7, isopropyl-D-1-thiogalactopyranoside (IPTG) was added to a final concentration of 0.2mM. Then the bacteria were grown at 16℃, 200rpm overnight to induce protein expression. The cells were collected by spinning down at 2200 rpm for 30 min. After resuspended with 20 mM Tris pH 7.5, 500 mM NaCl, resuspensions were homogenized with EmulsiFlex-C3. After spin down, the supernatant was collected and applied to HisTrap FF column (GE LifeSciences) and Resource S column (GE LifeSciences). The proteins are purified with AKTA Purifier system (GE Healthcare).

### NMR Protein purification

Uniformly ^15^N-labeled protein was expressed in Escherichia coli BL21(DE3) cells using minimal medium supplemented with ^15^NH_4_Cl as the sole nitrogen source. The recombinant plasmid encoding the target protein with an N-terminal His-tag was transformed into E. coli and grown overnight at 37 °C in LB medium containing 100 µg/mL ampicillin. The overnight culture was used to inoculate 1 L of M9 minimal medium containing 1 g/L ^15^NH_4_Cl (Cambridge Isotope Laboratories) and 4 g/L glucose. Cells were grown at 37 °C until the optical density at 600 nm (OD_600_) reached 0.8, at which point protein expression was induced by the addition of 1 mM isopropyl β-D-1-thiogalactopyranoside (IPTG). The culture was incubated for an additional 16 h at 37 °C before harvesting by centrifugation at 5,000 × g for 15 min. The cell pellet was resuspended in lysis buffer (50 mM Tris-HCl, pH 8.0, 300 mM NaCl, 10 mM imidazole) and lysed by sonication on ice. The proteins were purified on nickel column. After dialyzing the eluted protein, the His tags were cleaved using thrombin, and the proteins were then further purified by size-exclusion chromatography.

### NMR measurements

For peptides binding experiments, a series of ^1^H-^15^N HSQC experiments were performed at 24 °C on a 600 MHz Bruker Avance Neo NMR spectrometer equipped with a cryoprobe. The uniformly ^15^N-labeled Talin-F2F3 (400 μM) was prepared in 20 mM HEPES / 100 mM NaCl / 2 mM DTT / 5% D_2_O (pH 7.0). Peptide-binding experiments were performed by titrating small aliquots of peptides (20 mM in 100% DMSO) into the Talin-F2F3 solution, yielding final peptide concentrations of 100, 200, 300, 400, 500, and 600 μM. As a control, equivalent volumes of 100% DMSO were titrated in the absence of peptide. Peak assignments in ^1^H-^15^N HSQC spectra were based on the previously assigned resonances of Talin-F2F3 (BMRB ID: 15792).

The net change in chemical shift of each cross peak from Talin-F2F3, Δδ, was calculated as follows:

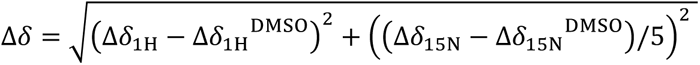

where Δδ_1H_ and Δδ_15N_ are the chemical shift changes in ^1^H and ^15^N dimensions for titration peptides in DMSO, respectively. Δδ_1H_^DMSO^ and Δδ_15N_^DMSO^ are the chemical shift changes in ^1^H and ^15^N dimensions for titration of DMSO, respectively. The spectra were processed using Topspin 4.2.0 software (Bruker) and analyzed using POKY.^43^

### CD spectroscopy

Circular dichroism (CD) spectra were acquired using a Jasco J-1100 spectrometer equipped with a temperature-controlled cuvette holder. Peptides were initially dissolved in DMSO, dried by vapor diffusion against 60% (w/v) NH₄NO₃ in DMSO, and subsequently resolubilized in 20 mM Tris-HCl (pH 7.5) and 100 mM NaCl to a final concentration of up to 0.4 mM. Measurements were performed using a 1 mm pathlength quartz cuvette. Spectra were collected from 190 to 250 nm at 0.1 nm intervals with a scan rate of 100 nm/min using three accumulations per scan. Baseline correction was performed by subtracting a buffer blank spectrum. Temperature interval scans were acquired from 20°C to 90°C in 10°C increments following temperature equilibration at each step.

### Cell permeability assay

The cell permeability assay was conducted as previously described.^21^ CHO-A5 cells were seeded in 6-well plates and then incubated with medium containing fluorophore-labeled peptides at various concentrations for 30 min at 37°C. The cells were washed twice with phosphate-buffered saline (PBS) and detached using 0.25% trypsin-EDTA. The detached cells were collected by centrifugation at 2,000 rpm for 5 min and resuspended in PBS and analyzed with an LSR flow cytometer (with a 525/50 filter and 488 nm laser), measuring 30,000 cells per sample.

### Fluorescent polarization (FP) assay and competition assays

The Fluorescent polarization assay and competition assay were carried out as described in previous studies.^21, 41^ Purified talin proteins at proper series concentrations (R7R8: up to 20 μM, THD: up to 40 μM) or buffer-only (20 mM Tris, pH 8.0, 100 mM NaCl, 2 mM DTT) were mixed with 20 nM FITC-labeled peptides (Genemed Synthesis, Inc.) in a buffer containing 20 mM Tris pH 8.0, 100 mM NaCl, 2 mM DTT, 0.5% Tween 20. After incubating for 3 minutes on ice, 20ul reaction mixture was added to individual wells of 384-well plate (Corning, 3573). Fluorescence polarization was then measured at room temperature using a Perkin Elmer Envision Plate Reader with FITC FP 480nm as excitation filter and a pair of FITC FP P-pol 535nm as emission filter. All the signals subtracted the buffer-only background signal were analyzed with Prism 10 and fitted to a single-site saturable binding model. For the competitive fluorescence polarization assay, various concentrations of unlabeled peptides were titrated into a solution containing 12µM R7R8 and 20nM FITC-TBS or a solution containing 3µM THD and 20nM FAM-b3. All the signals were subtracted buffer-only background signal and normalized based on the no competition data point. The competitive binding of unlabeled peptide to protein, leading to dissociation of FITC-labeled peptide, was indicated by a decrease in fluorescence polarization signals.

### Crystallization and structure determination

Purified R7R8 protein without tags was concentrated to 13.5 mg/ml a buffer containing 20 mM Tris pH 8.0, 80 mM NaCl and 2 mM dithiothreitol (DTT). A mixture of R7R8 and stapled peptides, S2-TBS (CPC Scientific) or S3-TBS (Vivitide), with a molar ratio 1:1.3 was subjected to crystallization, which was carried out at room temperature using the hanging-drop vapor diffusion technique. R7R8:S2-TBS crystals were harvested from a well solution containing 0.1 M Imidazole 8.0, 23% (w/v) polyethylene glycol (PEG) 8000, and 0.1 M NaCl. R7R8:S3-TBS crystals were harvested from a well solution containing 0.1 M Imidazole 8.0, 30% (w/v) polyethylene glycol (PEG) 8000, and 0.15 M NaCl. X-ray diffraction data were collected at the NSLS-II AMX and FMX beamlines of Brookhaven National Laboratory. The crystal structures were solved by molecular replacement using the structure of RIAM TBS in complex with talin R7R8 domains (PDB:4W8P) as a model and refined by Phenix and Coot.^44, 45^ The atomic coordinates and structure factors have been deposited in the Protein Data Bank with accession number 37TO and 37TQ for S2-TBS:R7R8 and S3-TBS:R7R8 complexes, respectively.

### GST pulldown

The pulldown was performed as described previously.^46, 47^ Purified his6-tagged talin rod protein, talin head protein and GST-tagged TBS1 proteins containing different mutations (wide type, I8L, T14W, L16W, G17W) were mixed in binding buffer (50 mM Tris, pH 7.5, 100 mM NaCl, 2 mM DTT) at a final concentration of 7.2uM, 5uM and 8.54uM individually. After settling down on ice for 15min, the protein mixtures were incubated with glutathione agarose beads (Invitrogen, G2879) which were pre-equilibrated with binding buffer. After rotating for 1h at 4°C, the supernatant was removed, and the beads were washed three times with binding buffer. The protein complex was eluted by incubating the beads with 50ul elution buffer (50 mM Tris pH 7.5, 100mMNaCl, 2mMDTT, 20mMreduced glutathione) at 4°C for 20min. The samples were resolved by SDS-PAGE and analyzed by Coomassie blue staining and Western Blotting. Anti-his (H1029-100UL) was used to detect his6-tagged proteins. Western blotting membranes were exposed by iBright 1500.

### Integrin activation assays

The integrin activation assay was performed as described previously.^48^ Briefly, CHO-A5 cells expressing αIIβ3 integrins were transfected with EGFP-C3-THD, wild-type or the W359A mutation, and incubated at 37°C for 24H. Cells were treated with TBS, S-TBS, S2-TBS and S3-TBS for 30min at 37°C. After washing with DPBS twice, the cells are detached with an enzyme-free cell dissociation solution (Sigma, c5914-100mL) and collected by gentle centrifugation. The cells were washed with DPBS and incubated with PAC-1 antibody (Invitrogen, MA5-28523, 1:50) in Tyrode’s buffer (136.9 mM NaCl, 10 mM HEPES, 5.5 mM Glucose, 11.9 mM NaHCO3, 2.7 mM KCl, 0.5 mM CaCl2, 1.5 mM MgCl2, 0.4 mM, NaH2PO4, pH 7.4) for 30 min at RT. After washing off the unbound antibody, the cells were incubated with Alexa 647 goat anti-mouse IgM (Jackson Immuno Research Labs, NC0401238, 1:400) in Tyrode’s buffer for 30min on ice in the dark. The cells were collected and resuspended with cold DPBS and quantified with a Symphony A5 analyzer (660/20 filter, 640 nm laser) using 10,000 cells per measurement. All the data were processed with FlowJo v9 and Prism 10.

### Quantification And Statistical Analysis

Crystallographic data processing and refinement were performed using COOT, PHENIX, and REFMAC. The Cell permeability data were processed using FlowJo and all the data normalized to the signal of the DMSO-treated or buffer-treated control samples. For the fluorescent polarization assay and competition assay, the buffer-only background signal was subtracted from all the data points and the data were processed using Prism 10. The results represent the mean of three independent experiments, with error bars indicating ± standard deviation. Statistical significance between groups was assessed using an unpaired two-tailed Student’s *t*-test.

### Computational mutagenesis

Structure-guided computational optimization of the pepetide was performed using FoldX v5.1. he crystal structure of the talin R7R8:TBS complex (PDB: 4W8P) and the NMR structure of the talin F3:TBS complex (PDB: 2MWN) were used as structural templates. For the THD:TBS complex, all 20 NMR conformers were extracted as individual PDB files. Each structure was processed using the RepairPDB function prior to mutational analysis. Point mutations across peptide residues Asp7-Ser25 were then generated using the BuildModel function. At each of the 19 peptide positions, the wild-type residue was individually replaced with each of the other 19 standard amino acids, resulting in 361 single substitutions for each talin complex. For the THD:TBS complex, mutagenesis calculations were performed for all 20 NMR conformers. For each substitution, the lowest ΔΔG among the 20 conformers was used for analysis. The THD and R7R8 datasets were combined by matching corresponding substitutions. A weighted score was calculated for each substitution using the equation:

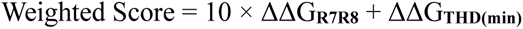

Substitutions were ranked according to their weighted scores. Structural models of selected TBS variants in complex with talin R7R8 and THD were generated using AlphaFold (AlphafoldServer.com)^49^.

### Gelatin degradation assay

The human breast cancer cell line BT-549 (Fox Chase Cell Culture Facility, Philadelphia, PA) was maintained in RPMI medium supplemented with 2 mM L-glutamine, 10% fetal bovine serum (FBS), and penicillin-streptomycin (50 U/mL penicillin and 50 μg/mL streptomycin, Corning). 24- or 96-well glass-bottom MatTek (MatTek Corporation) dishes were coated with fluorescently labeled gelatin as previously described.^50^ Briefly, glass-bottom MatTek dishes were first cleaned with 1 N HCl for 10 min and rinsed thoroughly with water. They were then coated with 50 µg/mL poly-L-lysine (Sigma-Aldrich) for 20 min, followed by several PBS washes. A 0.2% fluorescent gelatin mixture was prepared by diluting a 2% stock and combining it with thawed dye-labeled gelatin aliquots at a 1:8 ratio for 10 min. After the coating was removed, the surface was crosslinked using 0.2% glutaraldehyde on ice for 15 min, then neutralized with 5 mg/mL sodium borohydride (Sigma-Aldrich) for 15 min. The dishes were washed with PBS, disinfected in 70% ethanol, and finally incubated with penicillin/streptomycin in PBS. Prepared dishes were stored at 4 °C in the dark until use. BT-549 breast cancer cells were plated at 60,000 cells per well in 24-well glass-bottom plates or 20,000 cells per well in 96-well plates. Cells were allowed to attach and degrade the gelatin matrix for 20 h. Following incubation, cells were gently rinsed twice with PBS at room temperature and fixed with 4% paraformaldehyde for 10 min. Images of matrix degradation (appearing as black areas) and nuclear staining were obtained using a Nikon Eclipse Ti2-E widefield microscope equipped with a pco.panda sCMOS camera and a Plan Apo Lambda 60×/1.4 NA objective. Z-stacks were collected with a step size of 0.25 µm. Image processing and quantification were carried out in Fiji using a custom macro. Z-stacks were converted to maximum-intensity projections, thresholded, and analyzed using the “Analyze Particles” tool to identify degradation regions. To correct for differences in cell density across images, the total degradation area was normalized to the number of nuclei stained with DAPI or DRAQ5; (Thermo Scientific) within each field of view.

