## Supplemental Figure 1 for "Engineering Stapled Peptide Inhibitors Reveals Design Principles for Targeting Talin-Induced Integrin Activation"

**Supplemental Figure 1. Statistical analysis of stapled peptide inhibition in R7R8:TBS and THD-D397R:β3 competition assays**

**R7R8:TBS inhibited by peptides**

|  |  | 0 | 3μM | 6μM | 12μM | 24μM | 48μM |
| --- | --- | --- | --- | --- | --- | --- | --- |
| S-TBS | mean±stdev% | 100.0±5.5 | 96.0±12.7 | 83.1±1.9 | 78.2±7.7 | 69.1±10.2 | 61.8±31.5 |
|  | P value compare with 0 |  | 0.6731 | 0.0546 | 0.0464 | 0.0272 | 0.1476 |
| S2-TBS | mean±stdev% | 100.0±14.6 | 86.0±4.6 | 82.1±5.4 | 70.1±12.1 | 40.0±8.7 | 23.6±7.0 |
|  | P value compare with 0 |  | 0.2094 | 0.2138 | 0.1906 | 0.0044 | 0.0081 |
| S3-TBS | mean±stdev% | 100.0±10.6 | 79.7±8.9 | 68.9±11.3 | 62.0±4.0 | 56.8±21.7 | 46.8±12.8 |
|  | P value compare with 0 |  | 0.1862 | 0.1317 | 0.0415 | 0.0241 | 0.0370 |
| S4E | mean±stdev% | 100.0±10.1 | 82.4±16.3 | 72.1±2.6 | 70.6±4.6 | 61.4±3.5 | 67.3±2.2 |
|  | P value compare with 0 |  | 0.0752 | 0.0258 | 0.0456 | 0.0346 | 0.0216 |

**THD-D397R:β3 inhibited by peptides**

|  |  | 0 | 1.5μM | 3μM | 6μM | 12μM | 24μM |
| --- | --- | --- | --- | --- | --- | --- | --- |
| wt-TBS | mean±stdev% | 100.0±11.4 | 95.8±9.0 | 86.6±4.3 | 76.8±11.3 | 73.9±19.0 | 49.9±6.3 |
|  | P value compare with 0 |  | 0. 6161 | 0. 0821 | 0. 0265 | 0. 0428 | 0. 0135 |
| S-TBS | mean±stdev% | 100.0±8.7 | 90.6±6.0 | 75.7±8.1 | 52.6±7.0 | 44.7±9.4 | 37.2±10.1 |
|  | P value compare with 0 |  | 0.3779 | 0.0029 | 0.0014 | 0.0039 | 0.0037 |
| S2-TBS | mean±stdev% | 100.0±7.0 | 118.5±22.3 | 95.6±7.4 | 70.9±12.4 | 40.5±11.8 | 34.3±12.0 |
|  | P value compare with 0 |  | 0. 1802 | 0. 6159 | 0. 1191 | 0. 0291 | 0. 0264 |
| S3-TBS | mean±stdev% | 100.0±11.8 | 123.5±10.1 | 104.7±7.3 | 64.1±10.8 | 70.2±3.4 | 39.2±7.1 |
|  | P value compare with 0 |  | 0.1679 | 0.4929 | 0.1086 | 0.0584 | 0.0037 |
| S4E | mean±stdev% | 100.0±4.4 | 120.5±16.5 | 101.6±8.7 | 85.6±11.9 | 78.5±3.9 | 67.0±10.3 |
|  | P value compare with 0 |  | 0.1273 | 0.8094 | 0.2122 | 0.0239 | 0.0111 |
