## Supplemental Table 1 for "Engineering Stapled Peptide Inhibitors Reveals Design Principles for Targeting Talin-Induced Integrin Activation"

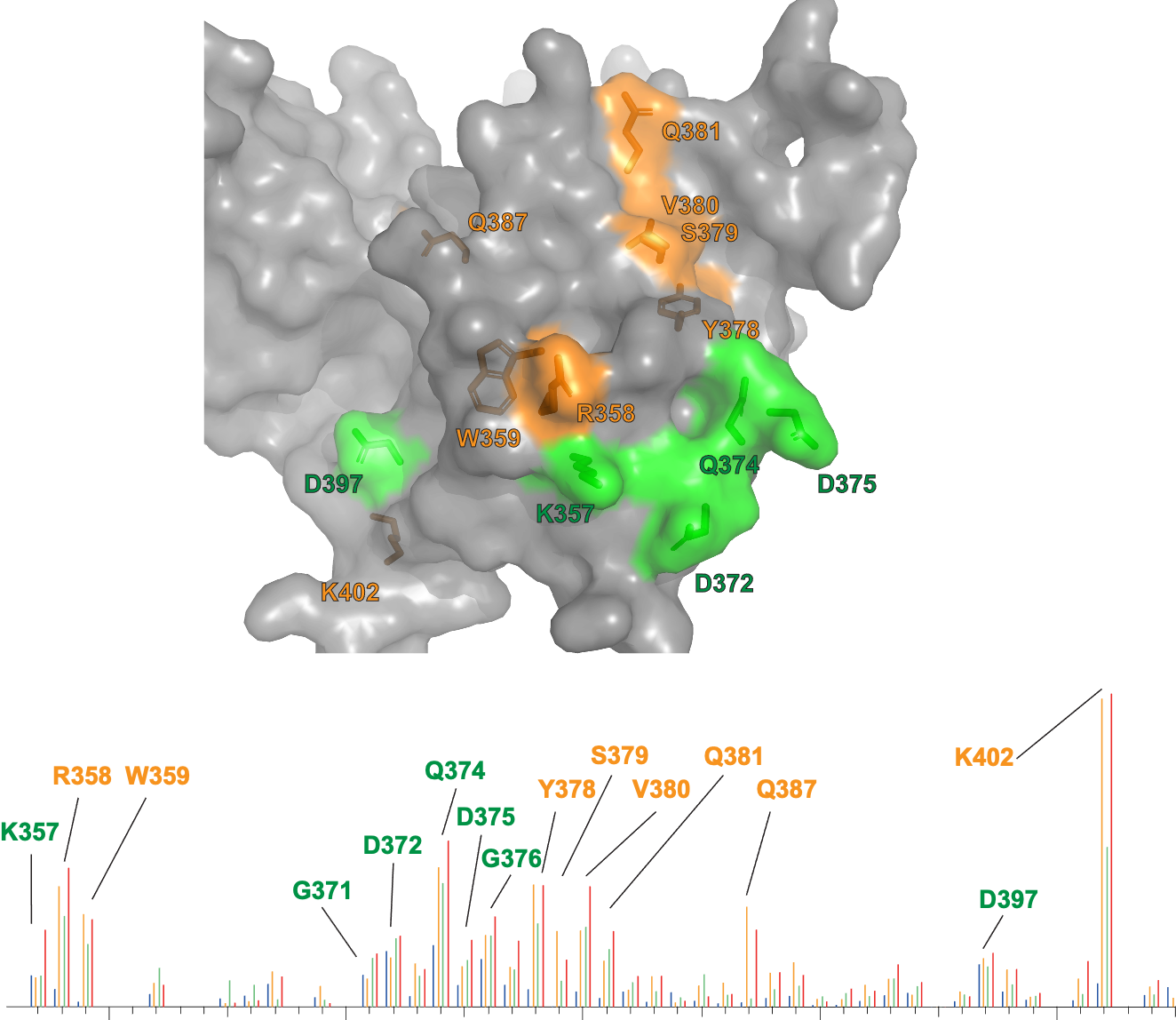

**Supplemental Figure 1. Stapled peptides enhance interaction with functionally important regions of talin F3.**

***Top***: mapped CSPs on the talin F3 surface induced by WT-TBS and stapled peptides. Conserved canonical interface residues are shown in green, whereas residues exhibiting enhanced perturbations with stapled peptides are shown in orange. ***Bottom***, residue-specific CSP comparison for WT-TBS (blue), S-TBS (yellow), S2-TBS (green), and S3-TBS (red) measured by ^1^H–^15^N HSQC NMR.
